# SARS-CoV-2 B.1.1.7 infection of Syrian hamster does not cause more severe disease and is protected by naturally acquired immunity

**DOI:** 10.1101/2021.04.02.438186

**Authors:** Ivette A. Nuñez, Christopher Z. Lien, Prabhuanand Selvaraj, Charles B. Stauft, Shufeng Liu, Matthew F. Starost, Tony T. Wang

## Abstract

Epidemiological studies have revealed the emergence of multiple SARS-CoV-2 variants of concern (VOC), including the lineage B.1.1.7 that is rapidly replacing old variants. The B.1.1.7 variant has been linked to increased morbidity rates, transmissibility, and potentially mortality (1). To assess viral fitness in vivo and to address whether the B.1.1.7 variant is capable of immune escape, we conducted infection and re-infection studies in naïve and convalescent Syrian hamsters (>10 months old). Hamsters infected by either a B.1.1.7 variant or a B.1 (G614) variant exhibited comparable viral loads and pathology. Convalescent hamsters that were previously infected by the original D614 variant were protected from disease following B.1.1.7 challenge with no observable clinical signs or lung pathology. Altogether, our study did not find that the B.1.1.7 variant significantly differs from the B.1 variant in pathogenicity in hamsters and that natural infection-induced immunity confers protection against a secondary challenge by the B1.1.7 variant.

## Introduction

Despite the proofreading activity provided by the 3’-5’ exonuclease activity of non-structural protein 14, Severe acute respiratory syndrome coronavirus 2 (SARS-CoV-2) has accumulated multiple mutations in its viral genome (2). Mutations occurring in the spike protein are of major concern due to the role of this glycoprotein in mediating virus entry and as the major target of neutralizing antibodies (3–6). In March of 2020, the D614G SARS-CoV-2 B.1 variant emerged and became the predominant strain of virus throughout Europe and the United States (7). Since late fall 2020, the emergent B.1.1.7 SARS-CoV-2 variant became predominant in the United Kingdom and contributed to the December surge in positive SARS-CoV-2 cases and the increase in hospitalization and death rates (8). Among the 8 mutations found in the spike protein of the B.1.1.7 variant, the N501Y mutation has been found to increase the binding affinity of spike protein to the human ACE2 receptor, resulting in an elevated transmissibility and monoclonal antibody resistance (9–11).

Here, we set out to address whether a prior exposure to an ancestral SARS-CoV-2 (D614) isolate would offer protection against re-infection by a B.1.1.7 variant. In addition, we investigated whether the B.1.1.7 variant displays fitness advantage over the G614 variant in the Syrian hamster model.

## Results

One group of convalescent (previously infected with Isolate USA-WA1/2020) and one group of naïve hamsters (n=12 per group, age paired) were intranasally inoculated with 10^4^ plaque forming units (PFU) of a U.S. B.1.1.7 variant (Isolate CA_CDC_5574/2020, termed CA B.1.1.7). A third group of 12 naïve hamsters (n=12) were similarly infected with a B.1 variant (Isolate New York-PV08410/2020, termed NY B.1). The circulating neutralizing antibody titers in the 12 convalescent hamsters ranged from 80 to 320 at the time of challenge. Nasal wash (NW) samples taken from infected hamsters on days 1, 2 and 3 post infection (PI) showed that all hamsters contained detectable sub-genomic RNA (sgRNA) levels in the nasal cavities on the first two days PI (Fig.1A). Three days following infection, viral sgRNA levels from nasal washes of convalescent hamsters declined to below the detection limit, indicating that convalescent hamsters quickly controlled B.1.1.7 replication. By contrast, naïve hamsters that were challenged with either the B.1.1.7 or B.1. variant showed 2-3 log_10_ higher levels of sgRNA compared to the convalescent group. Furthermore, naïve hamsters that were challenged with the B.1.1.7 variant displayed overall higher levels of sgRNA in NW samples (statistically significant at the day 2 PI) than those of the NY B.1 infected group (p=0.0008, 0.01 respectively). The same NW samples were also quantified for infectivity by an 50% tissue culture infectious dose (TCID_50_) endpoint dilution assay (Fig. 1B). Only 3 out of 12 NW samples from convalescent hamsters exhibited detectable levels of infectious virus at the day 1 PI, dropping to below detection at the day 2 PI (Fig. 1B). Surprisingly, NW samples from NY B.1 infected hamsters contained significantly higher TCID_50_ values than those from CA B.1.1.7 infected hamsters on days 2 and 3 PI (p=0.0109, 0.0016 respectively, Fig. 1B).

**Fig. 1.**
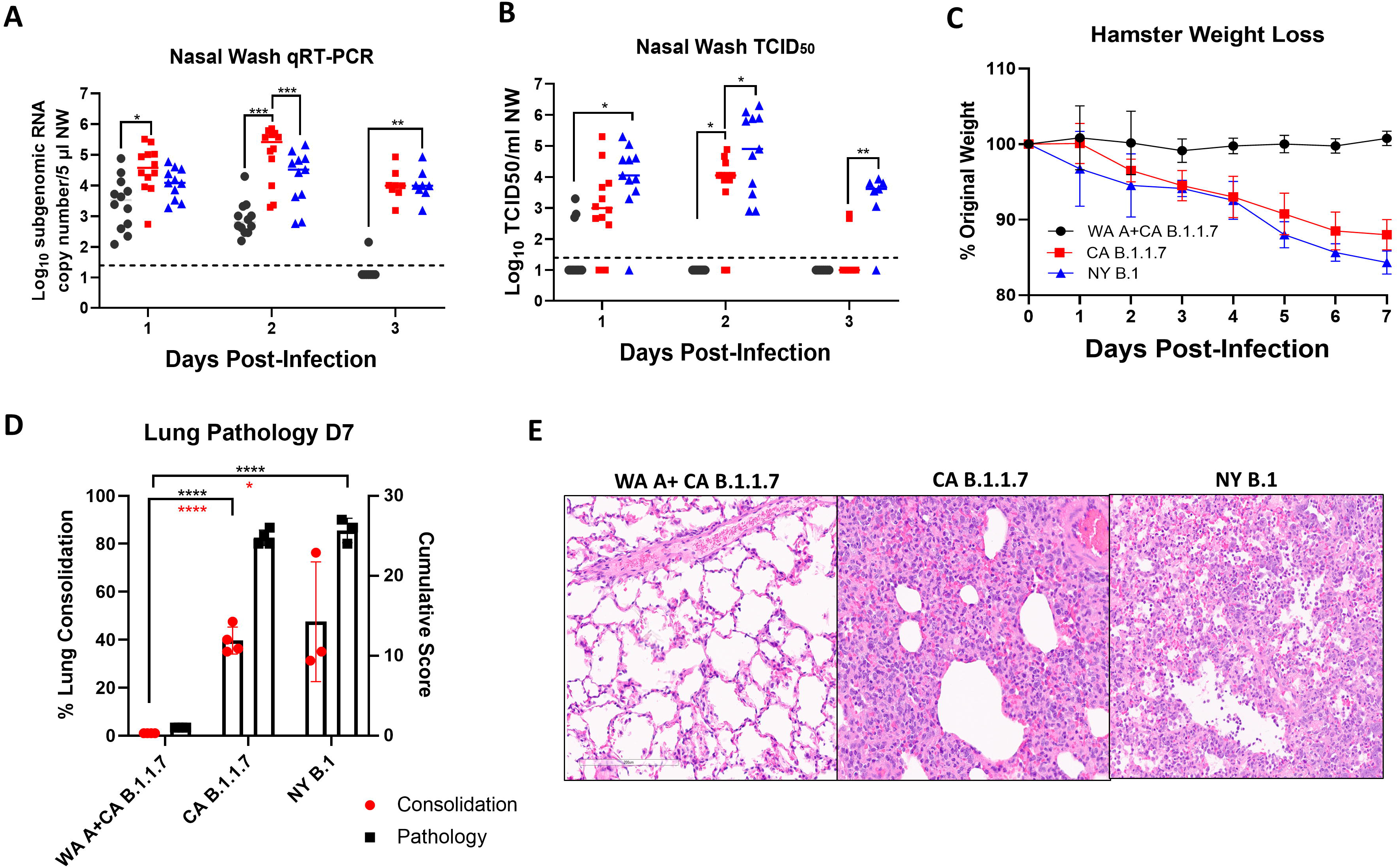
Convalescent hamsters are protected against severe disease from B.1.1.7 challenge. (*A*) Viral replication in nasal wash samples taken from hamsters on days 1-3 post-infection. (*B*) Shedding of infectious virus in nasal wash samples determined by an TCID_50_ assay. (*C*) Average weight loss was calculated based on initial weight taken on day 0 (day of infection). (*D*) Average pathology and consolidation scores from lung samples harvested at 7 days PI. (*E*) Representative images of hematoxylin and eosin staining of WA A+ CA B.1.1.7 (convalescent group), CA B.1.1.7 and NY B.1 infected animals. Data points represent values from single samples; bars represent means and standard deviations (STD).

Convalescent hamsters experienced no weight loss compared to the naïve infected groups over the course of seven days PI (Fig.1C). Both the CA B.1.1.7 and NY B.1 infected animals lost 10-15% of body weight by day 7 PI. At day 7 PI, infected hamster lungs displayed pathology including alveolar wall thickening, airway infiltrates, perivascular edema and hyperplasia (Fig. 1D-E). However, convalescent hamsters who were subsequently re-infected did not show these signature pathologies in the lungs (Fig. 1E). The cumulative pathology score and percentage of lung consolidation in the CA B.1.1.7 and NY B.1 naïve groups had no significant difference (Fig. 1D), but both had significantly higher scores than those convalescent hamsters (Fig. 1D). Therefore, although B.1.1.7 SARS-CoV-2 variant replication was higher in the nasal cavity on days 1 and 2 PI compared to the NY B.1 variant measured by sgRNA, the associated pathology was not more severe in the lung.

To account for the discrepancy between sgRNA titers and TCID_50_ titers in NW samples, a second study using ten male Syrian hamsters was subsequently performed. This study included two convalescent CA B.1.1.7 infected hamsters, four naïve NY B.1 and four naive CA B.1.1.7 infected Syrian hamsters aged >10 months. Again, NW samples of the CA B.1.1.7 infected hamsters show overall higher levels of sgRNA, but slightly lower levels of infectious virus than those of NY B.1 infected animals (Fig. 2A&B). Tissue samples taken from convalescent hamsters at 4 days PI revealed 3-4 log_10_ lower levels of sgRNA in all five lobes of the lung, trachea and nasal turbinate (NT) in comparison to those from naïve hamsters challenged with either the CA B.1.1.7 variant or the NY B.1 variant (Fig. 2C). Lung and NT homogenates were also titrated by plaque forming assay in Vero E6 cells (Fig. 2D). Homogenates taken from convalescent hamsters contained no detectable infectious virus at 4 days PI. Interestingly, NY B.1 infected hamsters showed significantly higher infectious viral titers than CA B.1.1.7 challenged hamsters (Fig. 2D).

**Fig. 2.**
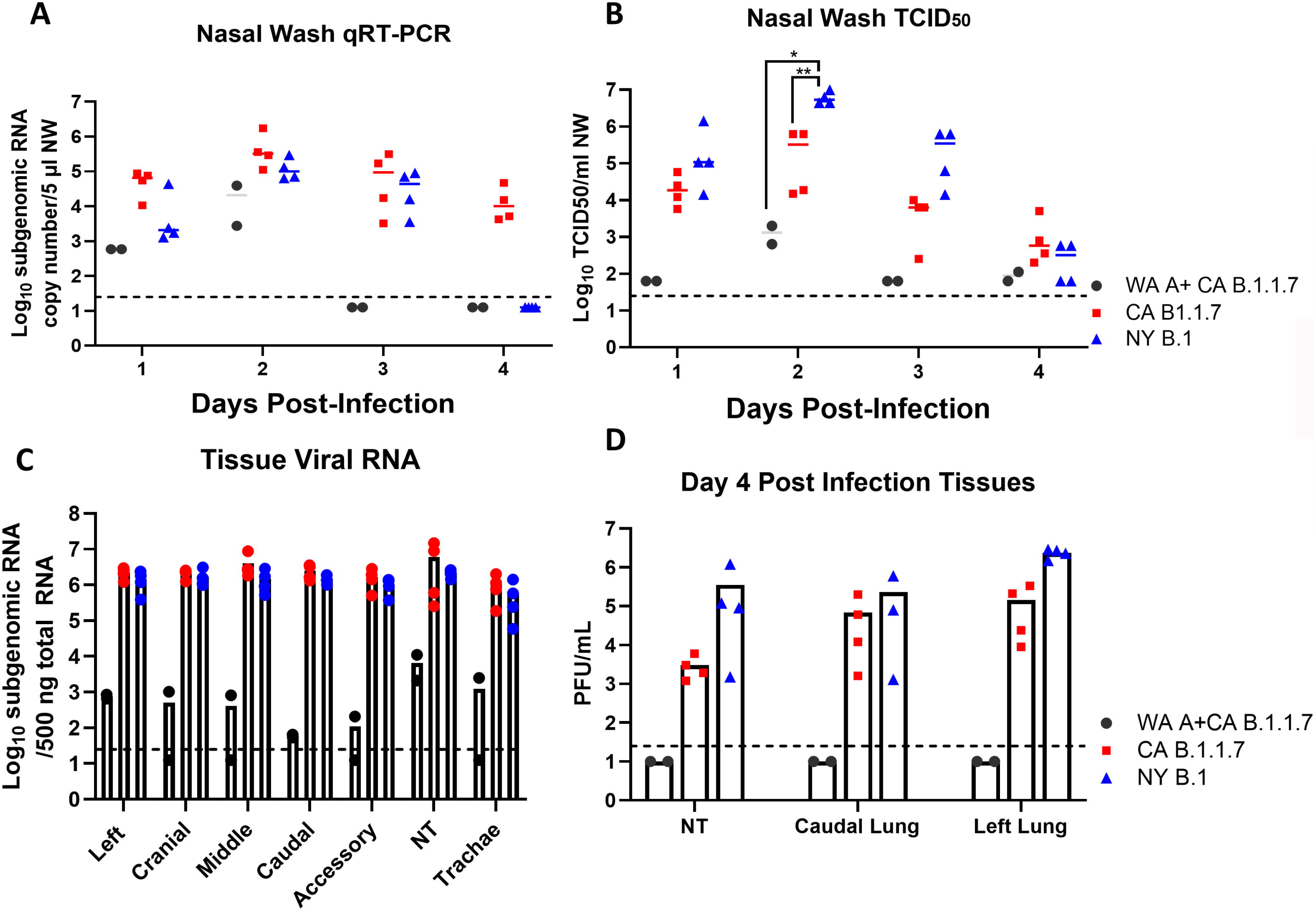
SARS-CoV-2 B.1.1.7 variant is not more pathogenic than the ancestral G614 variant. (*A*) Viral RNA was isolated from nasal wash samples taken from hamsters on days 4 PI. (*B*) TCID_50_ values of the same samples in A. (*C*) Lungs, NT and trachea tissues were collected 4 days post infection in 10 hamsters, sgRNA was measured via qRT-PCR. (*D*) NT and Lungs were tested for infectious virus by plaque assay. Dots represent single samples; bars represent means and standard deviations.

## Discussion

Altogether, this study finds that 1) convalescent hamsters are protected against interstitial pneumonia following B.1.1.7 variant challenge and that the B.1.1.7 variant of SARS-CoV-2 virus does not appear to be more pathogenic in adult male Syrian hamsters when compared to the B.1 variant (D614G). Differences in viral fitness may exist between the B.1.1.7 and B.1 variants in human beings, however, this was not observed in Syrian hamsters which may not be the ideal model to assess that possibility. This study did find an increase B.1.1.7 viral replication concomitant with reduced shedding of infectious virus in NW samples. Such a discrepancy of viral replication and infectious particles may be attributed to a disadvantaged B.1.1.7 plaquing effect in Vero E6 cells which has been previously mentioned (12) which would lead to undercounting of B.1.1.7 variant in Vero E6 cells. To determine whether the B.1.1.7 variant undergoes deleterious changes in hamsters that would alter the overall infectivity of the virus, we also performed next-generation sequencing of 16 hamster NW samples (8 from B.1.1.7 infected and 8 from B.1. infected hamsters). All sequences remain >99% identity with the sequence of the input virus. Notably, both the sgRNA and infectious titers of NW samples from the B.1.1.7 infected hamsters are higher than those of the NY B.1 infected hamsters at the Day 4 PI. This finding may suggest a prolonged viral shedding of the B.1.1.7, which has been hypothesized to increase the transmissibility of the B.1.1.7 strain in the human population (11, 13). However, this study did not find any increase in pathogenicity of the B.1.1.7 variant in hamsters. The significance of the increased replication of B.1.1.7 variant in the nasal cavity warrants further exploration through transmission studies.

## Materials and Methods

### Virus and cell culture

SARS-CoV-2 Isolate hCoV-19/USA/CA_CDC_5574/2020 and SARS-CoV-2/human/USA/NY-PV08410/2020 were propagated in Vero E6 cells to generate working virus stocks with infectious titers of 4.7×10^6^ pfu/ml and 1.8×10^7^ pfu/ml, respectively, and sequenced to confirm genotypes.

### SARS-CoV-2 pseudovirus production and neutralization assay

Procedures as described previously (14, 15).

### Hamster Challenge Experiments

Procedures as described previously (14, 15). For challenge studies, aged (10-12 months old) Syrian hamsters were anesthetized with 3-5% Isoflurane. Intranasal inoculation was done with 10^4^ pfu/hamster of SARS-CoV-2 in 50 μl volume dropwise into the nostrils. Nasal washes were collected by pipetting ~200 μl sterile phosphate buffered saline into one nostril. Male hamsters (n=12) were divided into three groups, the first group contained hamsters who were previously infected with the USA-WA1/2020 variant (lineage A, GISAID clade S) which was circulating in Washington State in early 2020. The second group of hamsters were immunologically naïve and were intranasally infected with the B.1.1.7 variant hCoV-19/USA/CA_CDC_5574/2020 (GISAID clade GR). The third group of hamsters were challenged with New York-PV08410/2020 (G614, B.1 lineage, GISAID clade GH). Following infection, hamsters were monitored for clinical signs and weight loss. Nasal wash samples taken on days 1, 2, and 3 post infection (PI) to test for sgRNA and TCID_50_. Seven-days following infection, a subset of hamsters was humanely euthanized and lungs for histopathology.

### RNA isolation and qRT-PCR

Procedures as described previously (14, 15).

### Histopathology Analyses

Procedures as described previously (14, 15).

### TCID_50_

Procedures as described previously (14, 15).

#### Plaque assay

Nasal wash samples were 10-fold serially diluted and added to a 6-well plate with Vero E6 cells. After 1 h the mixture was removed and replenished with Tragacanth gum overlay (final concentration 0.3%). Cells were incubated at 37°C and 5% CO_2_ for 2 days, then fixed with 4% paraformaldehyde, followed by staining of cells with 0.1% crystal violet in 20% methanol for 5-10 minutes.

### Statistical analysis

Standard unpaired T.test was used to calculate statistical significance through GraphPad Prism (8.4.2) software for Windows, GraphPad Software, San Diego, California USA, www.graphpad.com.

## Acknowledgments

We would like to thank the White Oak FDA Animal Program Staff and Veterinarians: John Dennis, Mario Hernandez, Mario Ortega, and Eric Nimako. We thank Ms. Katherine Shea for assistance with scanning pathology slides. The following reagents were deposited by the Centers for Diseases Control and Prevention and obtained through Biodefense and Emerging Infections Research Resources Repository, NIAID, NIH: SARS-Related Coronavirus 2, Isolate USA-WA1/2020, NR-52281; Isolate USA/CA_CDC_5574/2020, NR-54011; Isolate New York-PV08410/2020, NR-53514. The work described in this manuscript was supported by US FDA intramural grant funds. The funders had no role in study design, data collection and analysis, decision to publish, or preparation of the manuscript. The content of this publication does not necessarily reflect the views or policies of the Department of Health and Human Services, nor does mention of trade names, commercial products, or organizations imply endorsement by the US Government.

